# Genotypic Sex Determination Systems could be Adaptations to Extreme Temperature Environments in Reptiles and to Endothermy in Mammals and Birds

**DOI:** 10.1101/2022.02.07.479281

**Authors:** Manuel Ferrando-Bernal, Óscar Lao

## Abstract

In some vertebrates species, environmental temperatures (TSD) determine their sex determination. In others, it is controlled by genomic mechanisms (GSD). One hypothesis suggests that GSD systems could have evolved in ectothermic TSD species to escape the geographical limitation imposed by environmental temperatures. Recently, it has been found that TSD reptiles species tend to breed at warmer temperatures than GSD species, especially in habitat with four months of warm temperatures. Here we obtain the pivotal temperature (the one that generates equal ratios of male and females) in 53 reptiles species (from four orders: sphenodontia, crocodilia, testudines and squamata) and compare it with the environmental temperature in the nest during the breeding season in 100 TSD reptiles species and 78 GSD reptiles species. Our results show that GSD species statistically breed in temperatures that would cause a sex bias if they had TSD systems, whereas species with TSD breed in a similar range of temepratrues to the pivotal ones. Additionally, we also found that the body temperature of more than 1,200 endothermic species statistically exceeds the pivotal temperature suggesting that GSD is necessary for endotherms to avoid sex bias that could lead to extinction. Finally, we observed that one of the 100 most invasive species worldwide, *Trachemys scripta elegans*, a turtle species with TSD, has never been able to establish in countries with less than four months of warm temperatures, confirming a restriction in the geographic range in TSD species caused by extreme temperatures. Altogether, these results suggest that GSD could have evolved as an adaptation to avoid the biased sex ratios that extreme temperature may cause in species with a TSD system.

## 1. Introduction

Vertebrates show diversity in their sex determination systems (SDS) [1]. In some groups, such as some reptiles and fish, sex determination is produced polyphenically. In such organims sex determination mainly depends on changes in environmental temperature during embryonic development, a SDS known as Temperature-Dependent Sex Determination system (TSD) [2]. In these organisms, temperature above and below certain thresholds increases the likelihood to produce only one of the two sexes, which in some cases may lead to an incressment on the sex bias that could lead to extinction [3]. TSD is not a monotypic character. Depending on at which temperature(s) the sex bias(es) is produced, TSD is classified in distinctive patterns [4]. Pattern I is characterised by warmer temperatures producing one sex and cooler temperatures producing the other. This pattern is further divided depending on which sex is produced at warmer temperatures. In Pattern Ia, females are produced at warmer temperatures [4], whereas males are produced at these temperatures in Pattern Ib [5]. In Pattern II, both warmer and cooler temperatures produce one sex, while intermediate temperatures produce the opposite sex [4]. Nevertheless, a more complex TSD pattern has been reported in a particular species [6], suggesting that other TSD patterns are possible in nature.

A priori, the dependence of a SDS strategy based on temperature should impose some strong constraints to the environments that a TSD-based species can colonise. In particular, highly extreme environmental temperature areas –like in the Equator and polar latitudes-should impose a geographical limitation on the expansion of TSD species. According with this hypothesis, sex biases have been observed several times in sea turtles, ectotherm species with TSD, in whose habitats environmental conditions occassionally produce extreme temperatures as consequence of the Climatic change [7:9] as well as in other reptile orders, such as Sphenodontia [10]

Interestingly, it has been shown that SDS distribution is highly associated with environmental temperature, where TSD species tend to inhabit warm areas compared to GSD species, which inhabit independently in both cold and warm areas. In particular, TSD reptile species tend to inhabit areas where the environmental temperature during their breeding season is around 25 degrees, which may be the optimal temperature for nesting, for the sex-determining and for the embryo development [11]. This suggests that a strategy to elude the TSD constraints when colonising extreme temperature environments is to decouple the SDS from environmental temperature. In some vertebrate groups -including all mammals and birds, some reptiles, all amphibians, and many fish-sex determination is controlled by a genotypic mechanism named Genotypic Sex Determination system (GSD), where sex chromosomes are the best-known example [12,13]. Several hypotheses have been proposed for explaining in a given species the presence and maintenance of one SDS or another [14]. In this context, GSD systems may allow escaping the geographical limitation imposed by the environmental temperatures, and it may explain why GSD groups tend to inhabit at a higher geographical range (for example, birds, mammals, or snakes [15]), or why they tend to inhabit colder environments compared to TSD species [11, 16]. Some investigators claimed that the adaptive evidence for a given SDS is more scarce and controversial for many species, such as turtles. However, it seems that GSD may have evolved in some ectothermic species, such as some terrestrial turtles, to survive during the increasing environmental temperatures of previous climate changes [17] and interestingly, it has also been observed that sex chromosomes help populations living under extreme temperatures in ectotherm species of fish and lizards [18:21] whereas TSD populations of the same species inhabit less extreme environmental areas.

SDS is a plastic system from an evolutionary point of view. There is consensus that TSD is the ancestral sex-determination system in vertebrates, with GSD having evolved convergently through different systems of chromosomal sex determination [12,17, 22; 23]. Evolutionary transitions from GSD to TSD and vice-versa, or even both systems coexisting in the same species -where extreme temperatures during embryonic development can force a GSD strategy in TSD based species (e.g. sex reversal), have been reported in different ectotherm species [11, 24, 25], and even may occur in one generation [26]. Given such plasticity, it is intriguing that mammals and birds do not shift from GSD to TSD. Some authors have suggested that the lack of genetic variation, or the genomic instability that would follow a loss of one of the sex chromosomes, prevents the transition from one system to the other [24]. However, in that case, it is not explained why this restriction only applies to birds and mammals. A more parsimonious hypothesis can be formulated by relating the temperature constraints imposed by TSD and the temperature of bird and mammal embryo development. This temperature is closely related to their capacity to keep their body temperature independent of the environment -a feature known as endothermy [27].

In the case of birds, the optimal incubation temperature values are restricted to a narrow range (i.e. the optimal temperature in passerine birds is from 36°C to 40°C [28]. Extreme incubation temperatures, both above and below the optimal incubation temperature, can be lethal to bird embryos [28] or can be selectively lethal for a particular sex (as observed in *Alectura lathami* species [29). This introduces in the latest case a sex bias in the offspring. Furthermore, it represents a waste of resources for the mother in both situations. In the case of mammals, they have a gestation of, at least, two-thirds of the embryonic development [30; 31]. This viviparous feature forces the embryo to initiate its development under the high and constant temperatures of their mothers. These temperatures range from 30-32 °C in egg-laying mammals (the echidnas (*Bradypus*) and the platypus (*Ornithorhynchus*)), to 35 °C in marsupial mammals and to 36-39 °C in most placental mammals [32]. Interestingly, GSD systems are related to viviparity and also to extreme temperature environments [16]. It has been suggested that viviparity reduces the influence of the environmental conditions by manipulating the offspring’s development [33]. This independence of the environmental temperatures in viviparous species may explain why GSD is necessarily related to viviparity [16].

Therefore, it can be hypotesized that any TSD species whose embryonic development temperature is out of the pivotal temperature range of its TSD system is going to have difficulties generating an equal sex ratio in their offspring or producing viable descendants. Consequently, the GSD would be a necessary constraint in endothermic organisms. An addendum to this hypothesis would be that the acquisition of endothermy would be a cul-de-sac situation from any GSD-to-TSD transition.

A series of testable predictions described in table 1 can be derived from the proposed hypothesis.

**Table 1.** Hypotheses and predicted outcomes from the dependence between temperature and SDS tested in this paper.

| **Hypothesis** | **Predictions** |
| --- | --- |
| **1. Some GSD are adaptations to produce 1:1 sex ratios on extreme environmental temperature** | The environmental temperature during the breeding season should be close to the pivotal temperature in TSD species. Contrary GSD species would be able to breed over or lower these pivotal temperatures |
| **2. Invasive TSD species would be more common in the Temperate areas** | TSD species must become invasive only in countries where its environmental temperature is not extreme, whereas GSD species could be able to become invasive in a wider geographical range |
| **3. GSD is a necessary constraint in mammals** | Mammal body temperature exceeds the upper pivotal Temperature of ectothermic TSD species |
| **4. GSD is a necessary constraint in birds** | Bird body temperature exceeds the upper pivotal Temperature of ectothermic TSD species |

In this study we tested these hypotheses using different publicly available datasets. Our results suggest that the high body temperature of mammals and birds as well as the low temperature to where many GSD species inhabit may have caused sex bias in the populations, forcing the selection on GSD system, to maintain equal sex ratios and avoiding extinction.

## 2. Materials and Methods

### Databases

#### Breeding season temperature of ectothermic TSD species

The environmental temperature during the breeding season in 100 TSD and 78 GSD species were collected from [11].

#### The pivotal temperatures in TSD species

As the pivotal temperature of the TSD of the ancestor(s) of mammals or birds is unknown, we used as proxy the data for the pivotal temperature of 53 present-day TSD-ectothermic species from four different orders: six crocodilia, two sphenodontia, four squamata and 40 testudines species (See Supplementary Table 1). As we saw that both the endothermic groups tend to have a higer body temperature compared to the pivotal temperature of the TSD species, we used, when more than one pivotal temperature exist for a particular species (for example, in the cases were both high and low temperature produce a particular sex and intermediated temperature produce the other sex) the upper pivotal temperature.

We also use the pivotal temperature to analyse if the environmental temperature in TSD species is ideal to produce similar sex ratios. For this analysis, in the case where more than one pivotal temperature exists for a particular species we used the lower pivotal temperature to minimise the chance for a statistical difference. For these analyses we used the pivotal temperature of 53 present-day species.

#### Geographic distribution of invasive species

Invasive species are one of the major threats to natural ecosystems and native species. The International Union for Conservation of Nature has elaborated a list with the top 100 of the worldwide invasive species. One of them is a TSD turtle, the *Trachemys scripta elegans* which has been traded worldwide. Due to its dependence on the development temperature to generate males and females, we thought that this species may have difficulties to develop stable populations (a requirement to become invasive) in environments where there are few months of warm environmental temperature. To test it we use the data from the Invasive Species Compendium at the CABI.org. We used the data about the countries (and in some cases their specific locations) where *T. scripta elegans* has been declared an invasive species. We also recover the locations for five countries where it has never been observed in the wild. In order to test if *T.scripta elegans* is able to reproduce and establish new populations in climates with at least four month of warm temperatures we compare the climate in the countries and specific areas where it has declared invasive.

#### The Köppen-Geiger climate classification

For the Koppen-Geiger classification we used the data about the climate by latitude and longitude from [34]. Available online in http://koeppen-geiger.vu-wien.ac.at/present.htm

#### Birds and mammals body temperatures

Data of the body temperature of 664 birds and nearly 600 mammal species were taken from [35].

### Statistical tests

#### Hypothesis 1 & 2. Comparison with the pivotal temperature and the environmental temperature during the breeding season

A previous study [11] suggested that GSD systems are more frequent in reptiles inhabiting colder environments than TSD systems, raising the hypothesis that GSD could be an adaptation to avoid any sex bias caused by cold temperatures. In this analysis we decided to test if TSD group breed in temperature closer to the pivotal temperature at which they are able to produce equal sex ratios. Whereas GSD species will breed at lower or higher environmental temperatures as they would not suffer any sex ratio as their sex detemrination does not depends on the environmental temperature. To confirm if TSD group breed in temperature closer to the pivotal temperature at which they are able to produce equal sex ratios, we recopilate the pivotal temperature at which equal sex ratios are produced in 53 TSD species, and conducted the Mann-Whitney-Wilcoxon test complemented in the R software, comparing these data with the environmental temperature during the breeding season in 100 TSD species and 78 GSD species. The data was obtained from [11], and for these analysis we excluded the data referred to suborden serpentes, as the sex chromosome in this group is old and no GSD-TSD transition has been observed since then.

It has been shown that many reptiles are able to minimise the effects of environmental extreme conditions, the best known example is just nesting, but also choosing a particular nesting substrate or nesting shapes. Some studies have proven that nest conditions help to maintain a more stable and higher temperature compared to the temperature outside the nest. For example, in [36] found that the core nest temperature is between 2.8 and 2.3 degrees above the air temperature (31.7°C in the nest at a depth of 35 cm and 28.9°C and 29.4°C in the air). In [37] found that the individuals of the species *Caretta caretta* only dug a nest when they found areas with an increased temperature between 2.05 and 3.5 degrees. In [38] suggest that in a 27°C air temperature, nest of *Chelonia mydas* and *Eretmochelys imbricata* has a temperature of 3.8 °C in a Caribbean beach, 29.4°C and 31.3°C in Cape Verde beaches and 28.5°C and 31.5°C in Ascension island (depending on the sand colour). To approximate to the real values of the temperature at which the embryos develop inside the nest, we sum 3.016 °C (the mean values of the differences among the air temperature and the nest temperature collected from the commented studies) to all the values about the environmental temperatures during the breeding season for the 100 TSD and 78 GSD ectothermic species, and repeat the comparison with the lower pivotal temperature in the 53 TSD species (Supplementary Table 1).

#### Hypothesis 3. TSD species confront difficulties to invade extreme environmental climates

The Köppen-Geiger climate classification is one of the most commonly used climate classification systems. It divides climates into five main climate groups based on the temperature and precipitation patterns. These five main groups are referred to by a capital letter: A (Tropical), B (Dry), C (Temperate), D (Continental) and E (Polar). For each group (except the Polar) there is a second letter assigned which sub-classifies them depending on the precipitation pattern. The system also classifies the climates following their temperature patterns (except for the Tropical group) which refers to the second letter in the Polar group and to the third letter in the Dry, Temperate and Continental groups. Regarding the temperature patterns, the Tropical climates have a mean temperature over 22 °C and we can find four groups As, Af, Am, and Aw. The Dry climates are classified as “h” or “k” if the mean annual temperature is over or below 18 °C, as are classified into hot or cold deserts (BWh and BWk, respectively) and hot and cold semi-arid climates (BSh and BSk, respectively). For the Temperate and Continental climates, the letter “a” is assigned if the temperature is over 22 °C in at least a month and there are four with a temperature higher than 10 °C (Cfa, Csa, Cwa, Dfa, Dsa and Dwa climates), the letter “b” if there are at least four month over 0 °C but none with a temperature higher than 22 °C (Cfb, Csb, Cwb, Dfb, Dsb and Dwd climates), the letter “c” in no month is over 22 °C and there are among one to three months over 10 °C (Cfc, Csc, Cwc, Dfc, Dsc and Dwc climates). For the Polar climates, ET refers to climates with at least one month with an average temperature between 0 °C and 10 °C and EF if there is no month with an average temperature over 0 °C.

Recently, Cornejo-Páramo, P. et al. 2020 [11] found that TSD species tend to breed in periods of warmer temperatures. Specifically TSD species breed during the warmer months, with a mean of 4 breeding months. This limitation during the breeding season should influence the invasiveness capacity for TSD species as may confront difficulties to generate equal sex ratios in extreme temperatrue environments. So, following the Köppen-Geiger classification, it may be expected that TSD groups are present in Tropical and Dry climates, as well as in the Temperate and Continental climates but with a lower frequency in that climates with few warm months (Cfc, Csc, Cwc, Dfc, Dsc and Dwc climates) and difficult to be found in Polar climates (mainly in EF). To test if the invasiveness capacity of TSD species is restricted in these climates with few warm months we conducted a Fisher exact test using the R software [54] with data about the climates at the countries where the presence of *Trachemys scripta elegans* is considered invasive and the climates at the five countries where it has never been able to establish.

For the countries where it is not clear the location of where *T. scripta elegans* has developed an invasive population we selected the main climate at the country (South Africa was rejected for this analysis as many climates can be found). In the five countries where *T. scripta elegans* never established populations we used all the climates that can be found on them (Supplementary Table 2).

#### Hypothesis 4 & 5. GSD is a necessary constraint in endothermic organisms

Endothermy is a feature shared among mammals and birds. Their increased body temperature makes them more independent from the environmental temperature than ectotherms. Moreover, mammal embryos develop inside the body of their mothers and the range of optimal temperature at which the bird embryos develop is a bit narrow and over the upper pivotal temperature in TSD species. All these could diminish the range of the temperature at which the mammals and birds embryos develop and could lead to a potential sex bias. These two groups show GSD systems and, under such hypotheses, no transitions from GSD to TSD should be allowed. To test this idea, we conducted the Mann-Whitney-Wilcoxon test complemented in the R software [39] comparing the pivotal temperature of the 53 TSD-reptiles species with the body temperature of the 596 mammal and 664 bird species [35] (see Materials & Methods databases) (Supplementary Table 3).

## 3. Results & Discussion

The relationship among environmental temperature, SDS, sexual reproduction and the capacity to colonize new environments, acting as an evolutionary force for deciding which SDS is used, has been previously proposed for reptiles [17]. Several hypotheses, as depicted in Table 1, derivate, extend, or can be interpreted as direct consequences of such dependency. In particular, in this study we extended and tested the hypotheses related to the impact of endothermy in a TSD context, as well as the constriction of TSD species to breed around the optimal temperature to produce equal ratios of males and females. Additionally, we studied to what extent the presence of currently invasive TSD species follow the expected geographic pattern according to the hypothesis that environmental temperature conditionates the capacity of colonisation of a TSD species.

### 3.1. Ectothermic-TSD species breed at temperature that maximize equal sex ratios

Recently, [11] it has been suggested that the evolution of SDS systems in reptiles is related to temperature environment rather than to environmental temperature fluctuations. In particular, the authors reported that TSD species were more frequent in warmer climates cand GSD more frequent in cooler ones. Evenmore, they found that TSD species tend to breed in the warmer months of their breeding season compared to GSD species. In our study we have assessed this hypothesis by comparing the data of the lower pivotal temperature in 53 TSD species from several orders with the environmental temperature during the breeding season in 100 TSD and 78 GSD species (Supplementary Table 1).

Under our hypothesis we expect that TSD breed at temperatures close to the pivotal one at which they produce similar ratios of males and females. We observed statistically significant differences (see Methods) in both cases (a p.value of 1.105178e-10 among the pivotal temperature in the 53 TSD species with the breeding temperature in 78 TSD species and a p.value of 5.199568e-14 among the pivotal temperature in the 53 TSD species with the temperature during the breeding season in the GSD species). These results may indicate that there is a extended sex bias in mainly all the TSD species analysed here. However, we add 3.016 degrees over the environmental temperature during the breeding season in both the TSD and GSD species, to maximise the real values at which these species may develop (See materials and methods) and repeated the comparisons. The new results show that the difference among the TSD species breeding temperatures and the range of pivotal temperatures of the 53 TSD species is not statistically significant (p.value of 0.05896048)(see Figure 1). Whereas, the difference in the temperatures during the breeding season of the GSD species remains significant compared to the range of the pivotal temperature of the 53 TSD species. (p.value of 1.588705e-06) (Figure 1). These results suggest that TSD species tend to breed at temperatures that maximize the 1:1 sex ratios, whereas GSD species breed in temperatures that may cause extreme sex ratios if they had TSD systems.

**Figure 1.**
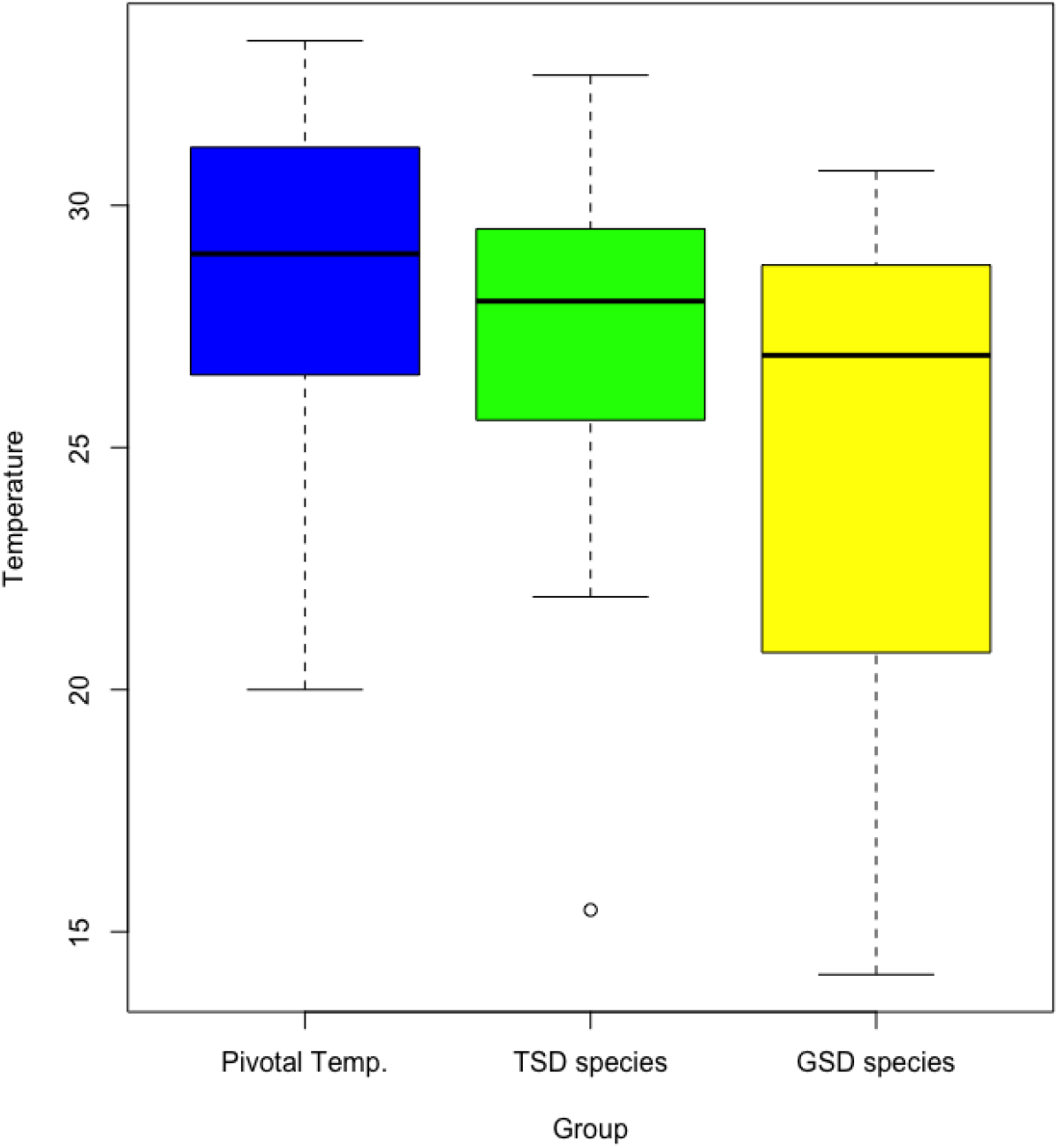
The Figure shows the range of temperatures during the breeding season in 78 GSD ectothermic species (in yellow, GSD species), in 100 TSD species (in green, TSD species) and the range of values of the pivotal temperature in 53 TSD species (in blue, Pivotal Temp.). The results show that the TSD species tend to breed in temperatures that produce equal sex ratios, whereas GSD species in temperatures that would cause sex bias if they had TSD systems.

Additionally, we tested if the pivotal temperatures are adapted to the environmental temperature during the breeding season in 46 of these 53 TSD species. We found a slight statistical correlation only among the lower pivotal temperature and the temperature during the breeding season (p.value 0.036, Adjusted R squared 0.075). These results suggest that the pivotal Temperature may be adapted to the breeding environmental temperature in each particular species, however this adaptation could not be so efficient. This is consistent with previous results where no changes in the pivotal temperature have been observed in different populations of *Chelonia mydas* that breed at different environmental breeding temperatures (40).

### 3.2 Invasive capacity of TSD species can be limited by temperature

Given our previous results, it should be expected that allochthonous TSD species should be only able to successfully colonise not temperature-extreme environments. We studied the invasive distribution of an ectothermic species with TSD, *Trachemys scripta elegans* [41]. Its natural habitat is the North Temperate zone, specifically between the southeast of the U.S.A. and the northeast of Mexico. It is one of the most worldwide traded turtle species [42] and one of the top-100 highlighted invasive species by the International Union for Conservation of Nature (IUCN) [43].

Due to its extensive pet trade and its posterior liberation, wild specimens have been reported in many countries (91.8% of the countries with data about the global distribution of *T.scripta* have reported seeing it wildly (Supplementary Table 2) [http://www.issg.org/index.html). Analysing data about the climates at the location where *Trachemys scripta elegans* has been declared invasive [http://www.issg.org/index.html], and the climates at the countries where it has never been found wildly, we found a statistically (Fisher Exact Test p.value = 0.00056) bias towards the climates with more than four months of warm average temperature (See **3.1. Ectothermic-TSD species tend to inhabit climates with at least four months of warm temperatures**). In contrast, *T. scripta* seem to be unable to establish populations in five countries where the climates have less than four months with an average of warm environmental temperature (Supplementary Table 2). These five countries are placed at high latitudes (Norway, Iceland, Greenland, Faroe islands and Estonia). These results suggests that *Trachemys scripta* has difficulties to develop stable populations and colonize some climates maybe due to the low temperatures in these areas during all the year likely lead to a male sex-bias.

Interestingly, in their natural areas, some populations of *Trahcemys scripta elegans* experience some sex ratio under temperature fluctuation, as they seem to increase the male rate as a consequence of the low temperature of the soil during the first months of the breeding season [44].

Overall, our results are consistent with the hypothesis that extreme temperatures difficult the geographical expansion of species with TSD. This could be a factor to take into account in conservationist projects to predict the likelihood that a TSD species, like *T. scripta*, may become invasive.

### 3.3 GSD is a necessary constraint in endothermic species

TSD seems to be the original SDS system in amniotes. Shifts from GSD to TSD have been observed several times in reptiles [14, 24, 25], whereas it has never been observed neither in mammals nor birds. The biological reason for this lack of GSD to TSD transitions is still debated. Different hypotheses have been proposed for explaining the lack of GSD to TSD in mammals and birds [24]. However, a more parsimonious explanation can be derived from the dependence between TSD and temperature. Since both mammals and birds are endothermic organisms, any transition from GSD to TSD in endothermic species should produce a sex bias. This would explain why there are no observed transitions from GSD to TSD in mammals or birds, whereas it is commonly observed in ectothermic species [14, 24, 25].

In order to test this hypothesis, we compared the body temperature of 596 different mammalian species [35] with the upper pivotal temperature over which there is a bias and most of the individuals likely develop as females in 53 TSD reptiles species (Supplementary Table 3). Mammals tend to have higher body temperatures than the upper pivotal temperature of the TSD species (Mann-Whitney-Wilcoxon test p-value = 1.993025e-31, Figure 2). Similar results were obtained when analysing 664 bird species (34) p-values (Mann-Whitney-Wilcoxon test p-value = 7.139647e-34). Therefore, all these results support the hypothesis that transitions from GSD to TSD in endotherms would lead to a important sex biase which may lead to extinction.

**Figure 2.**
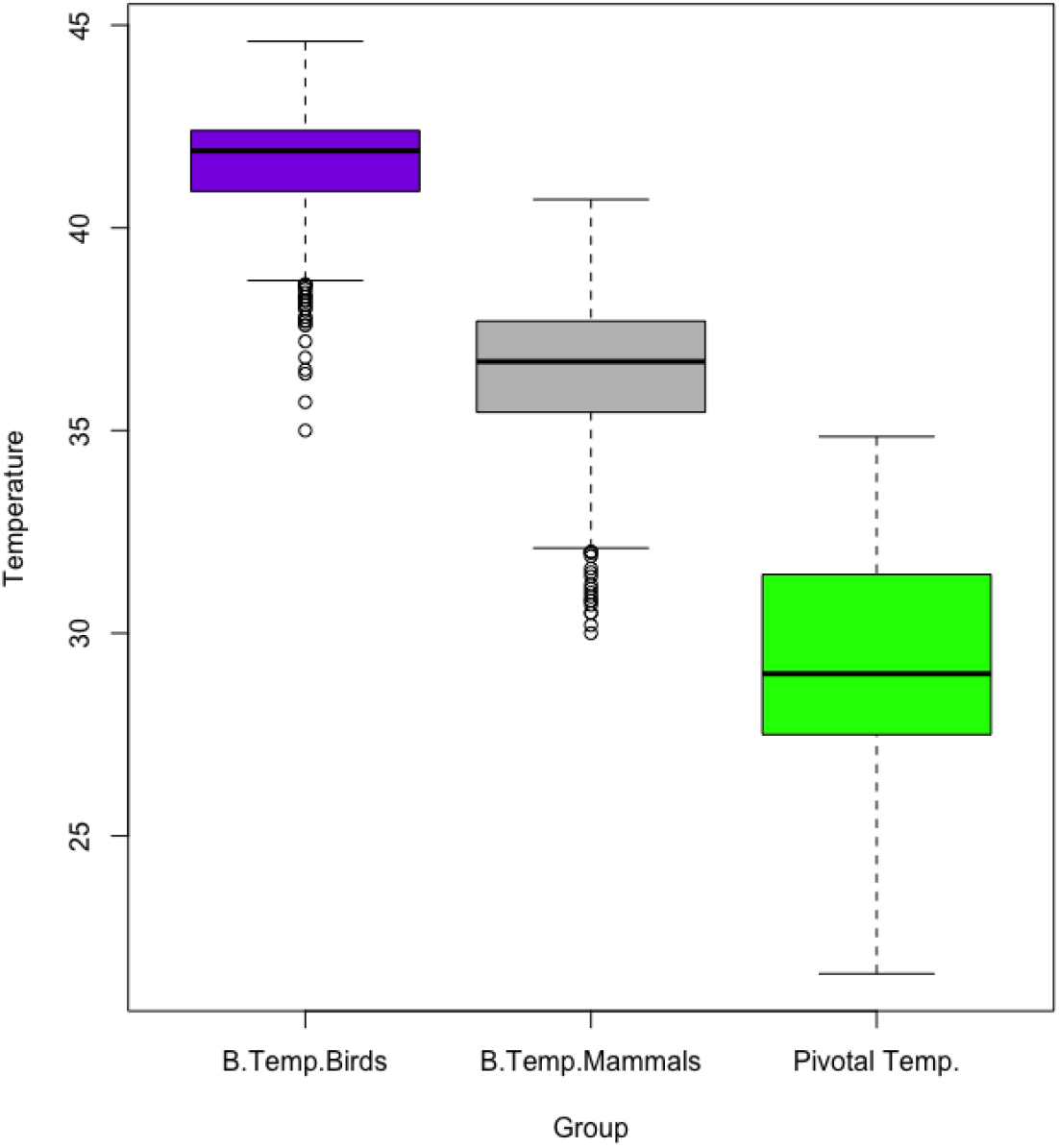
Box plot of the body temperature for 596 mammal species (grey, B.Temp. Mammals), 664 bird species (purple, B.Temp. Birds) and the upper pivotal temperature for female sex bias of 53 TSD ectotherm species (green, Pivotal Temp.). Differences are highly significant under a Mann-Whitney-Wilcoxon test, between endotherms and the upper pivotal temperature of ectotherms (p value of 1.993025e-31 between mammal body temperature and the ivotal temperature in reptiles; and 7.139647e-34 between bird body temperature and the pivotal temperature in reptiles).

## 4. Conclusions

Our results are consistent with the hypothesis that GSD systems such as sex chromosomes, could be adaptation to maintain equal sex ratios under specific situations. Specifically, where the embryonic development of the whole population is under or below the pivotal range, leading to a detrimental sex bias. In this study we observed that TSD species tend to breed inside the range of temperatures that allow to produce similar sex ratios in the offspring. However, GSD species breed at a statistical lower temperature values, likely leading to a sex bias. GSD system allow to maintain equal sex ratios as they do not depend on the temperature to produce different sexes in the offspring anymore. Additionally, we observed that a TSD worldwide traded species is unable to maintain wild populations in cold climates but has become invasive in countries where the temperature range allows it to incubate some nests over 25 °C and others in lower temperatures. Finally, we observed that both mammals and birds develop at high temperatures compared to the pivotal range in 53 TSD species, likely suggesting that any transition from GSD to TSD in these groups would lead to a sex homogenisation in the species which may lead to extinction, explaining why TSD systems has never been observed in these endothermic groups.

## References

1. Capel, B. Vertebrate sex determination: evolutionary plasticity of a fundamental switch. Nat. Rev. Genet. 2017, 18, 675–689.

2. Bull, J.J. Sex determination in reptiles. Q. Rev. Biol. 1980. 55, 3–21.

3. Mrosovsky, M.; Pierau, C. Transitional range of temperature, pivotal temperatures and thermosensitive stages for sex determination in reptiles. Amphibia-Reptilia. 1991, 12, 169–179.

4. Ewer, M.A.; Nelson, C.E. Sex Determination in Turtles: Diverse Patterns and Some Possible Adaptive Values. Cop. 1991, 1, 50–69.

5. Mitchell, N.J.; Nelson, N.J.; Cree, A.; Pledger, S.; Keall S.N.; Daugherty, C.H. Support for a rare pattern of temperature-dependent sex determination in archaic reptiles: evidence from two species of tuatara (Sphenodon). Front. Zool. 2006, 3, 9.

6. Inamdar Doddamani, L.S.; Vani, V.; Seshagiri, P.B. A tropical oviparous lizard, Calotes versicolor, exhibiting a potentially novel FMFM pattern of temperature-dependent sex determination. J. Exp. Zool. A. Ecol. Genet. Physiol. 2012, 317, 32–46.

7. Jensen, M.P.; Allen, C.D.; Eguchi, T.; Bell, I.P.; LaCasella, E. L.; Hilton, W.A. et al. Environmental warning and feminization of one of the largest sea turtle populations in the world. Curr. Biol. 2018, 28, 154–159.

8. Esteban, N.; Laloë, J.O.; Kiggen, F.S.P.L.; Ubels, S.M.; Becking, L.E.; Meesters, E.H.; Berkel, J.; Hays, G.C.; Christianen, M.J.A. Optimism for mitigation of climate warming impacts for sea turtles through nest shading and relocation. Sci. Rep. 2018, 8, 17625.

9. Laloë, J.O.; Esteban, N.; Berkel, J.; Hays, G.C. Sand temperatures for nesting sea turtles in the Caribbean: Implications for hatchling sex ratios in the face of climate change. J. Exp.Mar.Biol.Ecol. 2016, 474, 92–99.

10. Nelson, N.J.; Thompson, M.B.; Pledger, S.; Keal, S.N.; Daugherty, S. C. Do TSD, sex ratios, and nest characteristics influence the vulnerability of tuatara to global warming? Int. Congr. Ser. 2004. 1275, 250–257.

11. Cornejo-Páramo, P., Lira-Noriega, A., Ramírez-Suástegui, C., Méndez-de-la-Cruz, F.R., Székely, T.; Urrutia, A.O.; Cortez, D. Sex determination systems in reptiles are related to ambient temperature but not to the level of climatic fluctuation. BMC Evol. Biol. 2020, 20, 103.

12. Matsubara, K.; Tarui, H.; Toriba, M.; Yamada, K.; Nishida-Umehara, K.A.; Matsuda, Y. Evidence for different origin of sex chromosomes in snakes, birds, and mammals and step-wise differentiation of snake sex chromosomes. Proc. Nalt. Acad. Sci. U. S. A. 2006, 103, 18190–5.

13. Abbot, J.K., Nordén, A.K.; Hansson, B. Sex chromosome evolution: historical insights and future perspectives. Proc. Biol. Sci. 2017, 284, 20162806.

14. Valenzuela, N. Evolution and maintenance of temperature-dependent sex determination. In Temperature Dependent Sex Determination in Vertebrates. N. Valenzuela, V.A. Lance. Eds. Smithsonian Books, 2004, pp. 131–147.

15. Roll, U.; Feldman, A.; Novosolov, M.; Allison, A.; Bauer, A.M.; Bernard, R.; Böhm, M.; Castro-Herrera, F.; Chirio, L.; Collen, B. et al. The global distribution of tetrapods reveals a need for targeted reptile conservation. Nat. Ecol. Evol. 1, 1677–1682 (2017).

16. Cornejo-Páramo, P.; Dissanayake, D.S.B.; Lira-Noriega, A.; Martínez-Pacheco, M.L.; Acosta, A.; Ramírez-Suástegui, C.; Méndez-de-la-Cruz, F.R.; Székely, T.; Urrutia, A.O.; Georges, A. et al. Viviparous Reptile Regarded to Have Temperature-Dependent Sex Determination Has Old XY Chromosomes. Genome Biol. Evol. 2020, 12, 924–930.

17. Valenzuela, N.; Adams, D.C. Chromosome number and sex determination coevolve in turtles. Evolution. 2011, 65, 1808–13.

18. Pen, I.; Ullet, T.; Feldmeyer, B.; Harts, A.; While, G.M.; Wapstra, E. Climate-driven population divergence in sex-determining system. Nature. 2010, 468, 436–8.

19. Conover, D.O.; Heins, S.W. Adaptive variation in environmental and genetic sex determination in a fish. Nature. 1987, 326, 496–498.

20. Yamahira, K.; Conover, D.O. Interpopulation Variability in Temperature-Dependent Sex Determination of the Tidewater Silverside Menidia peninsulae (Pisces: Atherinidae). Cop. 2003, 1, 155–159.

21. Dufy, T.A., Hice, L.A. & Conover, D.O. Pattern and scale of geographic variation in environmental sex determination in the Atlantic silverside, Menidia menidia. Evolution. 2015, 69, 2187–95.

22. Vallender, E.; Lahn, B.T. Multiple independent origins of sex chromosomes in amniotes. PNAS. 2006, 103, 18031–18032.

23. Pokorná, M.J.; Kratochvil, L. What was the ancestral sex-determining mechanism in amniotes vertebrates? Biol. Rev. 2014, 91, 1–12.

24. Bachtrog, D. et al. Sex determination: Why so many ways of doing it? PLoS Biol. 2014, 12.

25. Quinn, A.E.; Georges, A.; Sarre, S.D.; Guarino, F.; Ezaz, T.; Marshall Graves, J.A. Temperature sex reversal implies sex gene dosage in a reptile. Science, 2007, 316, 411.

26. Holleley, C. E.; O’Meally, D.; Sarre, S.D.; Marshall Graves, J.A.; Ezaz, T.; Matsubara, K.; Azad, B.; Zhang, X.; Georges, A. Sex reversal triggers the rapid transition from genetic to temperature-dependent sex. Nature, 2015, 523, 79–82.

27. Hillenius, W.J.; Ruben, J.A. The Evolution of Endothermy in Terrestrial Vertebrates: Who? When? Why? Physiol. Biochem. Zool. 2004, 77, 1019–1042.

28. DuRant, S. E.; Hopkins, W.A.; Hepp, G.R.; Walters, J.R. Ecological, evolutionary, and conservation implications of incubation temperature-dependent phenotypes in birds. Biol. Rev. Camb. Philos. Soc. 2013, 88: 499–509.

29. Eiby, Y.A.; Wilmer, J.W.; Booth, D.T. Temperature-dependent sex-biased embryo mortality in a bird. Porc. Biol. Sci. 2008, 275, 2703–2706.

30. Ferrer, K.; Schultz, J.A.; Zeller, U. Comparative anatomy of neonates of the three major mammalian groups (monotremes, marsupials, placentals) and implications for the ancestral mammalian neonate morphotype. J. Anat. 2017, 231, 798.822.

31. Enjapoori, A.J.; Grant, T.R.; Nicol, S.C.; Lefèvre, C.M.; Nicholas, K.R.; Sharp, J.A. Monotreme lactation protein is highly expressed in monotreme milk and provides antimicrobial protection. Genome Biol. Evol. 22, 2754–73.

32. Ruben, J. THE EVOLUTION OF ENDOTHERMY IN MAMMALS AND BIRDS: From Physiology to Fossils. Annu. Rev. Physiol. 1995, 57, 69–95.

33. Ma, L.; Buckley, L.B.; Huey, R.B.; Du, W-G. A global test of the cold-climate hypothesis for the evolution of viviparity of squamate reptiles. Glob. Ecol. Biogreogr. 2018, 27, 679–689.

34. Kottek, M., J. Grieser, C. Beck, B. Rudolf, and F. Rubel, 2006: World Map of the Köppen-Geiger climate classification updated. Meteorol. Z., 15, 259–263. DOI:10.1127/0941-2948/2006/0130.

35. Clarke, A.; Rothery, P. Scaling the body temperature in mammals and birds. Funct. Ecol. 2008, 22, 58–67.

36. Mhady, A.; Ahmed, A.F.A.; Idris, M.Y.M.; Mohamed, I.M.A.; Samie, M.A.A.; Mayor, M.A.A.; Emgower, S.A.B.; Said, R.E.M. The effects of nesting ground temperatures on incubation and hatchability of loggerhead turtle Caretta caretta inhabiting the Mediterranean Sea Coast, Libya. Egypt. J. Aquat. Biol. 2020. 24, 5: 111–114.

37. Stoneburner, D.L. & Richardson, J.I., Observations on the role of temperature in j loggerhead turtle nest site selection. Copeia, 1981. 1: 238–241.

38. Esteban, N.; Laloë, J-O.; Mortimer, J.A.; Guzman, A.N.; Hays, G.C. Male hatchling production in sea turtles from one of the world’s largest marine protected areas, the Chagos Archipelago. Sci. Rep. 6: 20339.

39. R Core Team. 2021. R: A language and environment for statistical computing. R Foundation for Statistical Computing, Vienna, Austria. URL http://www.R-project.org/.

40. Tilley, D.; Ball, S.; Ellick, J.; Godley, B.J.; Weber, N.; Weber, S.B.; Broderick, A.C. No evidence of fine scale thermal adaptation in green turtles. J. Exp. Mar. Bio. Ecol. 2019. 514-515: 110–117.

41. Wibbels, T.; Cowan, J.; LeBoeuf, R. Temperature-dependent sex determination in the red-eared slider turtle, Trachemys scripta. J. Exp. Zool. 1998, 281, 409–416.

42. Cadi, A.; Joly, P. Impact of the introduction of the red-eared slider (Trachemys scripta elegans) on survival rates of the European pond turtle (Emys orbicularis). Biodiversity and Conservation. 2004, 13, 2511–2518.

43. Lowe, S.; Browne, M.; Boudjelas, S.; De Poorter, M. 100 of the World’s Worst Invasive Alien Species A selection from the Global Invasive Species Database. Published by The Invasive Species Specialist Group (ISSG) a specialist group of the Species Survival Commission (SSC) of the World Conservation Union (IUCN). Aliens. 2000, 12. Updated and reprinted version: November 2004.

44. Tucker, J.K.; Dolan, C.R.; Lamer, J.T.; Dustman, E.A. Climatic Warning, Sex Ratios, and Red-Eared Sliders (*Trachemys scripta elegans*) in Illinois. Chelonian Conserv. Biol. 2008, 7, 1: 60–69.

